# Increased Telomere Mobility in Progeria is Restored by Isoprenylcysteine Carboxyl Methyltransferase Inhibition

**DOI:** 10.64898/2026.04.25.720781

**Authors:** Gabriella Gagliano, Alex Raterink, Xijie Yang, Martin Bergö, Anna-Karin Gustavsson

**Affiliations:** Department of Chemistry, Rice University, Houston, TX, 77005, United States; Smalley-Curl Institute, Rice University, Houston, TX, 77005, United States; Applied Physics Program, Rice University, Houston, TX, 77005, United States; Department of Radiation Physics, University of Texas MD Anderson Cancer Center, Houston, TX, 77030, United States; Systems, Synthetic, and Physical Biology Program, Rice University, Houston, TX 77005, United States; Department of Medicine, Karolinska Institutet, Huddinge, Sweden; Department of BioSciences, Rice University, Houston, TX, 77005, United States; Department of Electrical and Computer Engineering, Rice University, Houston, TX, 77005, United States; Center for Nanoscale Imaging Sciences, Rice University, Houston, TX, 77005, United States; Department of Cancer Biology, University of Texas MD Anderson Cancer Center, Houston, TX, 77030, United States

## Abstract

Hutchinson-Gilford Progeria Syndrome (HGPS) is a genetic disease characterized by the accumulation of progerin, a mutant form of lamin A, at the nuclear envelope. Progerin disrupts the stability of the nuclear lamina, leading to genome instability and accelerated aging phenotypes. While structural nuclear defects are well-documented, the impact of progerin on real-time chromatin dynamics and the ability of current therapeutics to rescue these dynamics remains poorly understood. In this work, we employ single-particle tracking to quantify telomere dynamics in HGPS patient fibroblasts. We demonstrate that HGPS cells exhibit significantly increased telomere dynamics, characterized by expanded scan areas, increased diffusion coefficients, and larger jump distances compared to healthy controls. We further evaluated the efficacy of two clinically relevant treatments, the farnesyltransferase inhibitor Lonafarnib and the ICMT inhibitor C75, to determine if emerging treatments can restore chromatin dynamics compared to healthy controls. Our results reveal that Lonafarnib partially rescues telomere dynamics, shifting chromatin motion back towards healthy control levels, and that C75 provides a complete rescue of the dynamics for all parameters quantified. These findings provide a quantitative framework for understanding how nuclear lamina mutations induce aberrant genome dynamics and the efficacy of HGPS therapies on restoring those dynamics.

## Introduction

Nuclear lamins play a crucial role in cell health by interacting closely with chromatin to support various vital cellular functions including cell cycle regulation, gene expression, and chromatin organization^1–4^. However, in Hutchinson-Gilford Progeria Syndrome (HGPS), a mutation to lamin A leads to significant nuclear abnormalities that disrupt mechanotransduction^5^ and genome dynamics^6–8^, causing shortened telomeres^9–13^ and accelerated aging^14–16^. HGPS is a genetic disorder most commonly caused by a single heterozygous *de novo* point mutation (c.1824 C>T) in LMNA, the gene which encodes the nuclear lamina protein prelamin A^17,18^. In healthy cells, prelamin A matures into lamin A, which localizes to the nuclear periphery and the nucleoplasm^19,20^ and is an important protein in the nuclear lamina meshwork. However, in HGPS, progerin, the mutated form of prelamin A, cannot be converted into mature lamin A, and accumulates at the nuclear membrane^14,21^, where it forms an abnormally organized lamina meshwork with enlarged inter-filament spacing^22^. Previous studies have shown that progerin stiffens the nuclear envelope, which in turn leads to a weakened nuclear force response^23–25^. The mislocalization of progerin to the nuclear periphery also results in a softening of chromatin within the nuclear interior^23^. Additionally, it is known that lamin A acts as an important linker for chromatin in the nucleoplasm and at the nuclear periphery, and without it, telomeres become untethered^26,27^. However, the effect of progerin on telomere dynamics has not been directly quantified in HGPS fibroblasts. Given that premature cellular senescence in HGPS results from progerin-induced telomere dysfunction^12,28^, understanding the connection between nuclear lamina defects and their effects on chromatin behavior is highly relevant for understanding of HGPS disease pathology and other genetic nuclear lamina-related diseases^29–31^.

Several treatment strategies have been proposed for HGPS, including pharmacological approaches to mitigate progerin toxicity^32^ and newly emerging *in vivo* genome editing approaches demonstrated in mouse models^33–35^. However, the only current FDA-approved treatment for HGPS is farnesyltransferase inhibitor (FTI) Lonafarnib^36,37^. Inhibition of protein farnesylation blocks the first post-translational modification step of progerin in which the cysteine residue in the CAAX motif is farnesylated by the enzyme farnesyltransferase^38,39^, thereby disrupting the formation of the mutated progerin protein and partially mislocalizing it away from the cell membrane where it normally accumulates^40–42^. Previous studies demonstrated moderate amelioration of HGPS disease phenotypes^43,44^ and lower mortality rates in both mouse models^45–48^ and humans^49–51^ after treatment with FTIs. However, treatment with FTIs have also been associated with several cellular side effects^52–54^.

Another promising treatment strategy for HGPS focuses on inhibiting the methylation of progerin during a later post-translational step in which the farnesylated cysteine residue is methylated by isoprenylcysteine carboxyl methyltransferase (ICMT)^55^. By targeting ICMT^56^, both by genetic inactivation of the ICMT gene and by pharmacological intervention with small molecule ICMT inhibitors, previous works have succeeded in decreasing the severity of HGPS-related defects and increasing survival in mouse models^57–59^.

Live-cell fluorescence imaging enables visualization of dynamic cellular processes in their native environments at high spatial and temporal resolution. Given that chromatin in the cell nucleus is highly organized and yet very dynamic, the sensitivity and specificity of live-cell fluorescence microscopy makes it an ideal choice for tracking the motion of chromatin at the nanoscale^60^. Previously, tracking of telomeres in lamin A knockout cells revealed that loss of lamin A increased chromatin dynamics in the nuclear interior^26,27,61^, suggesting that chromatin behavior is highly dependent on lamin A regulation. Given that in HGPS, lamin A is partially mutated into progerin, we hypothesized that telomeres in HGPS cells would be similarly affected due to the reduction of tethering by healthy lamin A. It has been previously shown that telomeres in progerin-expressing cells demonstrate aberrant and increased contact with the nuclear lamina^62,63^, and that chromatin-associated nucleolar proteins in progerin-expressing and HGPS patient cells showed cell-type-dependent results, with both increased and decreased ensemble-averaged mean squared displacements (MSDs) depending on the cell type^64^. However, while chromatin motion has been studied in engineered progerin-expressing models, or by tagging chromatin-bound proteins as proxies, high resolution single-particle tracking of telomere-specific dynamics has yet to be performed in primary HGPS patient fibroblasts. Additionally, treatment with FTI and ICMT inhibitors has been shown to redistribute progerin from the nuclear rim and back into the nucleoplasm^59,65–67^, but the efficacy of these treatments for restoring chromatin dynamics in patient cells has yet to be quantified.

To address this knowledge gap and gain a more complete picture of how chromatin dynamics is affected in HGPS, we used single-particle tracking to quantify telomere motion in HGPS patient fibroblasts and the effect of two different progerin-targeting drugs on telomere behavior to identify if pharmaceuticals could restore chromatin dynamics to healthy control levels (Fig. 1). We found that telomeres in HGPS cells exhibit markedly increased dynamics, characterized by increased scan areas, faster diffusion, and larger jump distances compared to healthy controls. Treatment with FTI Lonafarnib^36,68–70^ partially rescued telomere behavior, with decreased average scan areas and diffusion coefficients, whereas treatment with ICMT inhibitor C75^65^ fully restored telomere dynamics to healthy levels. Here, we provide a direct, quantitative view of telomere dynamics at the nanoscale to elucidate how nuclear lamina-related mutations alter chromatin motion, providing insights into cellular senescence and disease mechanisms in HGPS and across the broader spectrum of laminopathies.

**Fig. 1.**
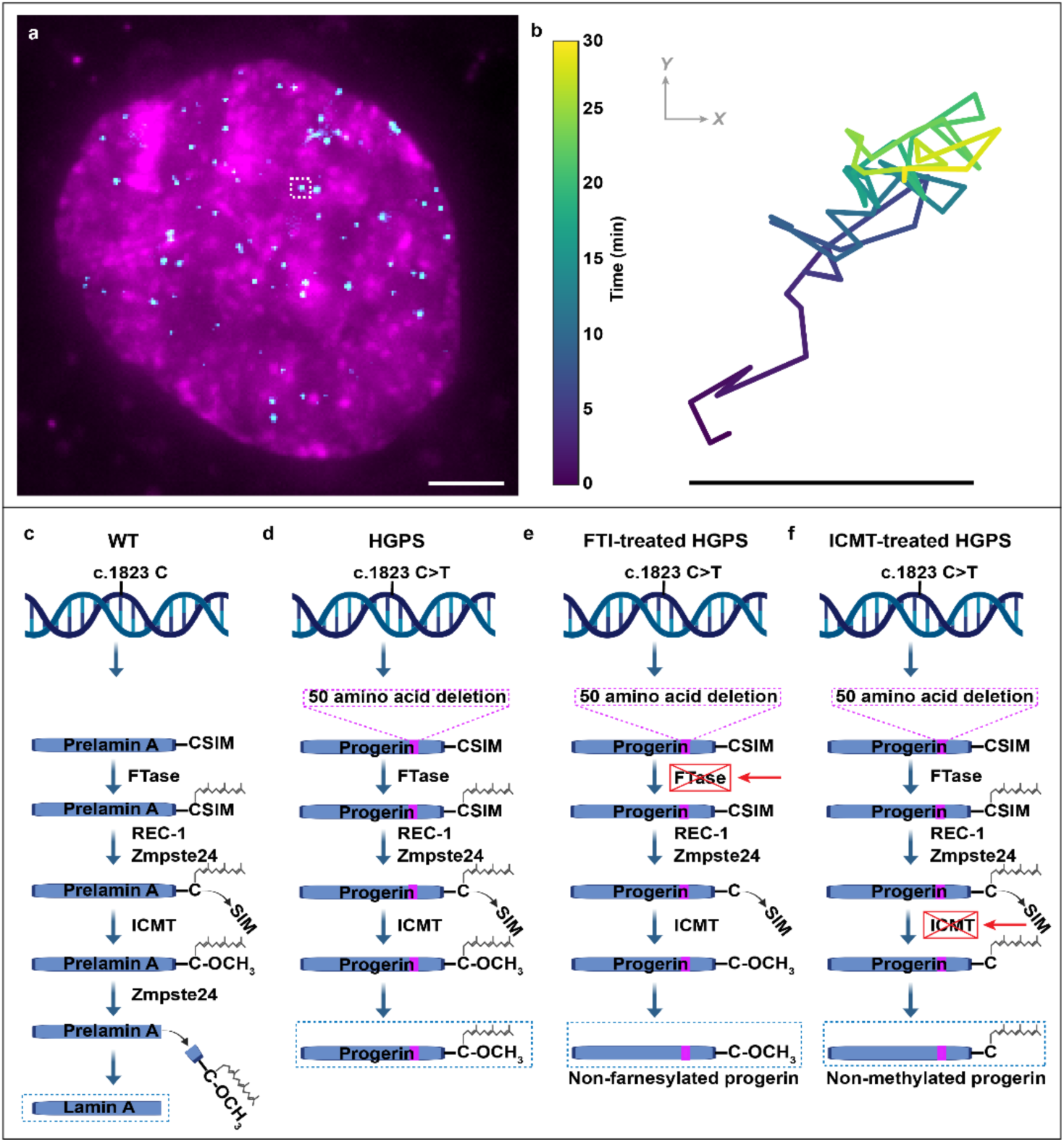
Live-cell telomere tracking and post-translational modification steps of lamin A in healthy cells and progerin in HGPS cells. **a** HGPS cell nucleus labeled with silicon rhodamine (SiR)-DNA nuclear stain in magenta and a maximum-intensity z-projection of telomeres tagged with fluorescent protein mStayGold in cyan. Scale bar 5 μm. **b** Example trajectory from **a**, demonstrating telomere tracking over 30 min. Scale bar 500 nm. **c** Post-translational modification steps of prelamin A. First, the cysteine residue is farnesylated by the enzyme farnesyltransferase (FTase). Next, the last three amino acids (SIM) are removed by the RAS-converting enzyme RCE1 or by zinc metalloproteinase ste24 (Zmpste24). Then, the farnesylated cysteine residue is methylated by isoprenylcysteine carboxyl methyltransferase (ICMT). Finally, the entire carboxyl section is cleaved off by the protease Zmpste24 and mature lamin A is formed. **d** Post-translational modification steps of progerin in HGPS. Mutation in the LMNA gene in exon 11 (c.1824 C>T) leads to an abnormal splicing site, resulting in a 50 amino acid deletion in the mutated prelamin A protein. This elimination occurs around the Zmpste24 cleavage site, resulting in a retention of the farnesylated and methylated C-terminal and the production of progerin. **e** Post-translational modification steps of progerin when treated with farnesyltransferase inhibitors (FTIs), preventing the farnesylation of the cysteine residue. **f** Post-translational modification steps of progerin when treated with ICMT inhibitors, preventing methylation of the cysteine residue.

## Results

### Telomeres in HGPS cells exhibit larger scan areas which are partially or fully restored by pharmacologic treatment

To quantify the effects of HGPS on chromatin dynamics, we performed single-particle tracking of telomeres in HGPS patient fibroblasts as well as in healthy wild-type (WT) fibroblasts from the father of the HGPS donor. Telomeres were labeled by transducing cells with a plasmid expressing the monomeric green fluorescent protein StayGold (mStayGold)^71,72^ fused to the telomeric repeat binding factor 2 (TRF2)^73,74^ (Supplementary Fig. 1). Individual trajectories of labeled telomeres were captured using highly inclined and laminated optical sheet (HILO) microscopy^75^ and tracked by acquiring z-stacks of telomere-labeled cells every 30 s for 45 min. From these trajectories the scan areas of telomeres in HGPS cells and healthy control cells were extracted and quantified (Fig. 2a-e). We found that in HGPS cells, telomeres scanned significantly larger areas as compared to healthy control cells (Fig. 2c-f, Supplementary Fig. 2), signifying a reduction of physical confinement.

**Fig. 2.**
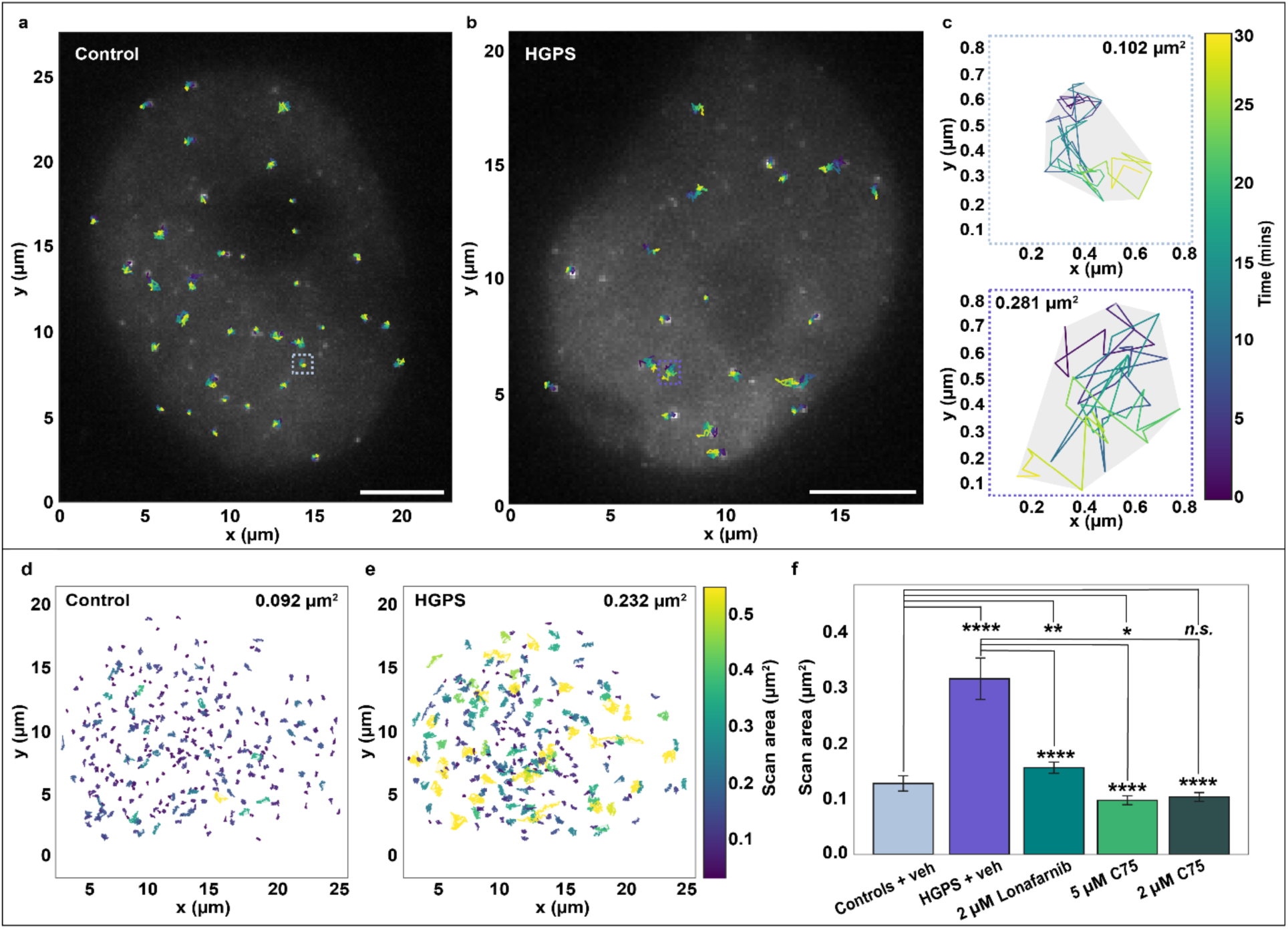
Telomere scan area analysis and response to FTI and ICMT inhibitors. **a** Representative example of a nucleus from the healthy control group and **b** the HGPS group with overlayed telomere trajectories tracked over 30 min. Scale bars 5 μm. **c** Selected telomere trajectories from the dashed boxes (top) for the healthy cell in panel **a**, and (bottom) for the HGPS cell in panel **b**, with scan areas indicated. **d** Telomere scan areas from 307 trajectories from 12 different cells in the control group and **e** telomere scan areas from 254 trajectories from 15 different cells in the HGPS group, colored by scan area, with mean scan areas indicated. **f** Bar chart showing the mean ± SEM of per-cell mean telomere scan areas for all five conditions. The per-cell mean values were pooled across the three replicate groups per condition. *n.s*.: p > 0.05, *: *p* < 0.05, **: *p* < 0.01, ****: *p* < 0.0001.

We next evaluated whether pharmacological interventions could rescue the increased HGPS telomere dynamics. HGPS cells were treated with either 2 µM of the FTI inhibitor Lonafarnib^36,68–70^ for 2 days, or 5 µM of the ICMT inhibitor C75^65,76^ for 20 days before transduction. C75 was also tested at a lower concentration and shorter duration of 2 µM for 10 days. Treatment with Lonafarnib partially restored telomere scan areas back to control levels (Fig. 2f, Supplementary Fig. 2). Notably, C75 had a more robust reduction in scan area compared to Lonafarnib, effectively restoring telomere scan areas back to baseline levels, or slightly below in the case of the 5 µM treatment condition (Fig. 2f, Supplementary Fig. 2). The performance of C75 in rescuing telomere dynamics suggests that it may be more efficient at redistributing progerin and establishing a re-confinement of chromatin territories. Treatment with C75 also influenced HGPS cell growth, where both concentrations outperformed Lonafarnib in stimulating cell proliferation (Supplementary Fig. 3).

**Fig. 3.**
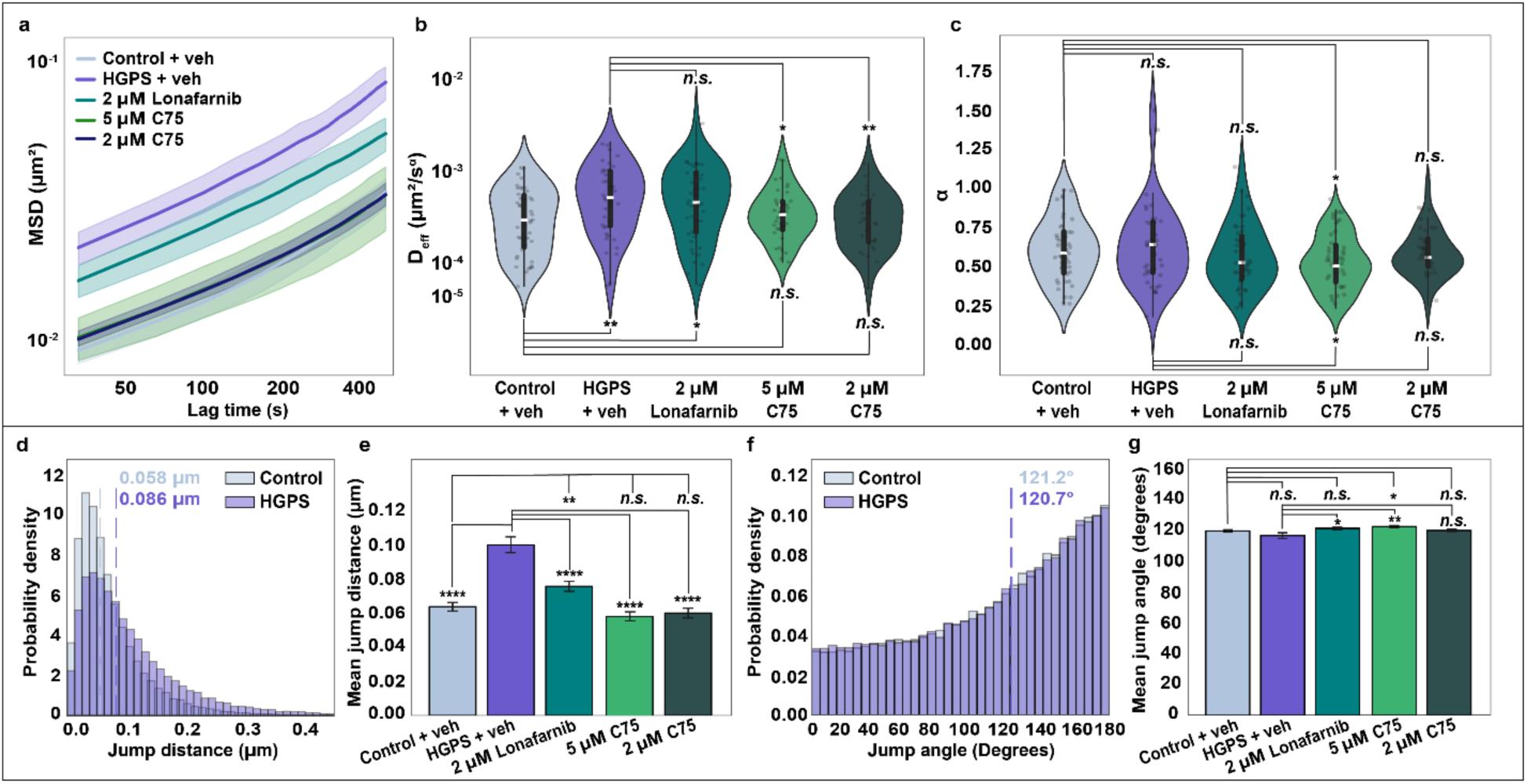
Telomere dynamics in HGPS and response to FTI and ICMT inhibitor. **a** MSD comparison between telomere trajectories in healthy control cells, HGPS cells, and HGPS cells treated with either FTI Lonafarnib or ICMT inhibitor C75. Data shown is the mean ± SEM of three technical replicates acquired across three different days. **b** Violin plots of effective diffusion coefficients (D_eff_) and **c** anomalous exponents (α) extracted from fits of the MSD data in **a**. The data are shown as cell medians of three technical replicates acquired across three different days. **d** Histograms comparing telomere jump distances in healthy control and HGPS cells. All individual jumps from all tracked telomeres were pooled across three technical replicates acquired on three separate days. Dashed vertical lines indicate the median jump distance for each group. **e** Bar plot showing the mean ± SEM of the per-cell median jump distance. The per-cell median values were pooled across the three replicate groups per condition. **f** Histograms of consecutive jump angles for telomeres in control and HGPS cells. Dashed vertical lines indicate the median jump distance for each group. All angles were pooled across the three technical replicates per condition. **g** Bar plot showing the mean ± SEM of the per-cell median jump angle. The per-cell median values were pooled across the three replicate groups per condition. *n.s*.: p > 0.05, *: *p* < 0.05, **: *p* < 0.01, ****: *p* < 0.0001.

### Telomeres in HGPS fibroblasts show increased effective diffusion coefficients and jump distances that are partially or fully restored by pharmacologic treatment

Next, for each trajectory, the 2D mean squared displacement (MSD), effective diffusion coefficient (D_eff_), anomalous exponent (α), jump distances, and jump angles were extracted.

Telomeres in HGPS cells were found to have increased MSDs (Fig. 3a, Supplementary Fig. 4) compared to healthy controls, as well as significantly increased effective diffusion coefficients (Fig. 3b, Supplementary Fig. 4), likely resulting from reduced chromatin tethering in the nuclear interior. The anomalous exponent did not significantly change in HGPS cells as compared to healthy control cells (Fig. 3c, Supplementary Fig. 4).

**Fig. 4.**
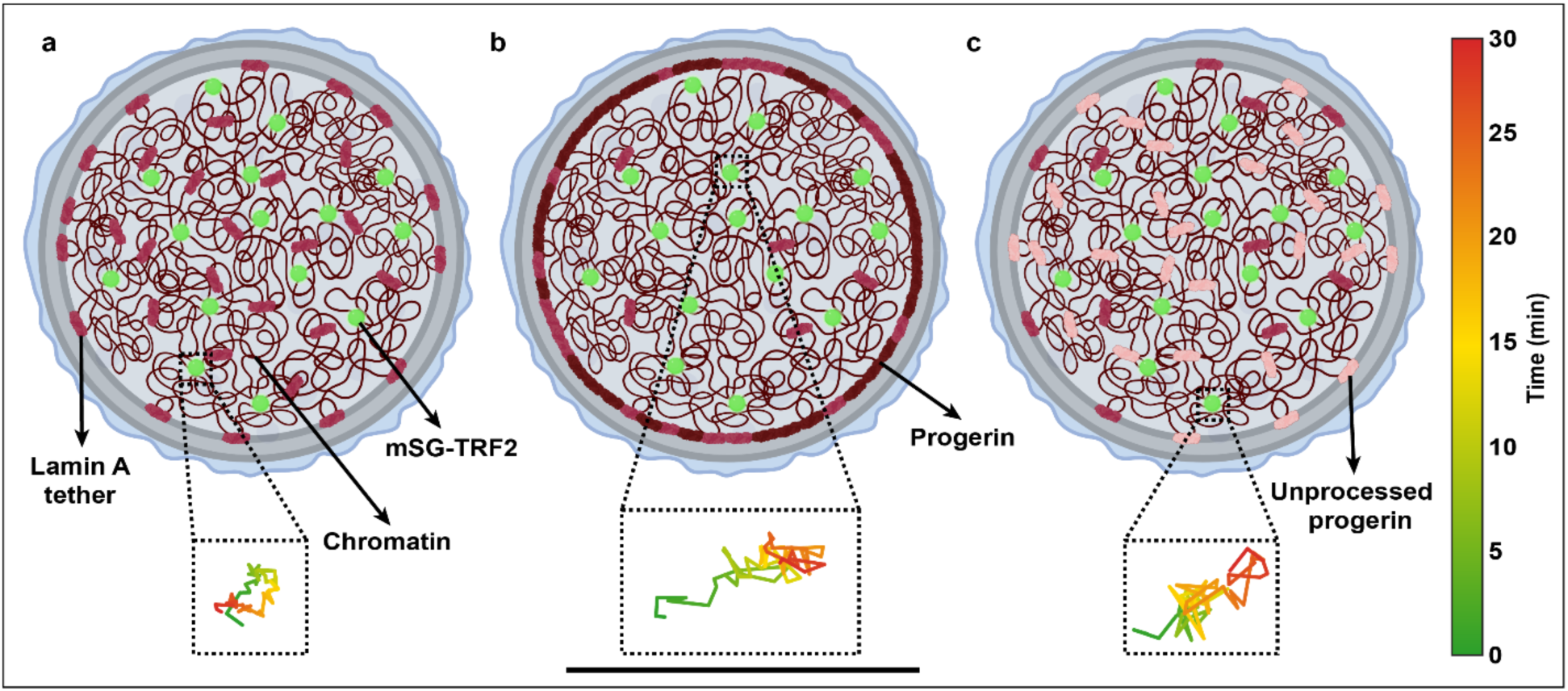
Proposed model for progerin-induced chromatin dynamics dysregulation. **a** Model of a healthy nucleus, where both nucleoplasmic lamin A and lamin A at the nuclear periphery help anchor chromatin. Fluorescently tagged telomeres are shown in green. Zoomed in inset shows a representative telomere trajectory as seen in healthy cells. **b** Model of a nucleus with HGPS. Progerin is accumulated at the nuclear envelope, resulting in reduced lamin A anchoring in the nuclear interior. Wild type lamin A still exists in the nuclear interior due to the heterozygous nature of the mutation. Zoomed in inset of a telomere trajectory showing increased telomere dynamics due to a reduction in lamin A anchoring. **c** Model of pharmacological rescue of telomere dynamics in an HGPS nucleus by either the farnesyltransferase inhibitor (FTI) Lonafarnib or the isoprenylcysteine carboxyl methyltransferase (ICMT) inhibitor C75. Unprocessed progerin, in which either the farnesylation or the methylation of the protein has been blocked, is redistributed off of the nuclear rim and back into the nucleoplasm where it works to anchor chromatin. Inset telomere trajectory shows a partial return to healthy dynamics. Scale bar 1 μm.

Next, cells were treated with either 2 µM Lonafarnib, 5 µM C75, or 2 µM C75. Treatment with Lonafarnib reduced the MSDs and effective diffusion coefficients of HGPS telomeres, although not to a significant degree (Fig. 3a,b, Supplementary Fig. 4). However, both treatments with C75 had a significant effect on telomere dynamics, returning both the MSDs and effective diffusion coefficients back to healthy control levels (Fig. 3a,b, Supplementary Fig. 4). The anomalous exponent did not show a significant change across any group except the 5 µM C75 group (Fig. 3c, Supplementary Fig. 4).

While FTI and ICMT treatments do not restore the 50-amino-acid deletion characteristic of progerin, our data suggests that the spatial redistribution of the mutant protein off of the nuclear rim and into the nucleoplasm by these drugs is sufficient to partially or fully rescue chromatin dynamics (Fig. 3). By chemically inhibiting these post-translational modifications (Fig. 1e,f), progerin is released back into the nucleoplasm, where it retains enough structural motifs to re-establish transient tethers with chromatin.

Next, we investigated the effects of HGPS and treatments on the frame-to-frame consecutive jump distances and jump angles. The jump distances of telomeres in HGPS nuclei were found to be significantly elevated as compared to healthy controls (Fig. 3d,e, Supplementary Fig. 5a), indicating that reduced tethering allows telomeres to take larger jumps, in agreement with the MSD analysis results. Treatment with both Lonafarnib and C75 significantly decreased the mean jump distance (Fig. 3e, Supplementary Fig. 5a) of telomeres. The most significant effects were seen after treatment with C75, which restored the mean jump distance to that of the healthy control. Consecutive jump angles were then calculated as the angle between successive displacement vectors. Jump angle analysis revealed that telomeres in both HGPS and control cells showed a 180° jump bias (Fig. 3f Supplementary Fig. 5b) demonstrating anti-persistent motion. Previous measurements have shown that chromatin motion is generally constrained^77–79^ with jump preferences towards 180°, or backward stepping^80^. After treatment with FTI and ICMT inhibitors, the jump angle showed small differences between groups (Fig. 3g, Supplementary Fig. 5b), consistent with the anomalous exponent results (Fig. 3c).

## Discussion

In this study we investigated the effects of HGPS on chromatin dynamics and evaluated the restorative potential of pharmacological interventions. Our data reveal that HGPS patient fibroblasts exhibit significantly increased telomere dynamics, evidenced by expanded scan areas, higher diffusion coefficients, and larger jump distances relative to healthy controls. This hyper-mobility may stem from the structural instability of the nuclear lamina, possibly due to gaps in the progerin meshwork^22^, and a reduction of intranuclear lamin A, which weakens the tethering of chromatin both to the nuclear periphery and in the nuclear interior (Fig. 4a,b). By disrupting these physical constraints, progerin effectively softens the nuclear interior.

To assess the clinical relevance of these findings, we tested the efficacy of the FDA-approved FTI Lonafarnib alongside the ICMT inhibitor C75. Our results demonstrate that Lonafarnib can partially rescue telomere dynamics, shifting chromatin behavior towards slower and more constrained motion characteristic of healthy cells^26,74,79,81^, possibly by restoring laminar anchors. Strikingly, we find that treatment with ICMT inhibitor C75 can fully restore chromatin dynamics to healthy cell levels for all parameters measured. The ability of some of these drugs to stabilize chromatin motion, even partially, suggests that inhibiting the post-translational processing of progerin can restore a degree of order to the genome.

As FTIs prevent farnesylation altogether, they also broadly impair the ability of other CAAX-containing proteins to localize to the cell membrane, including nuclear envelope proteins such as lamin B1 and the RAS family proteins which are critical for regulating cell proliferation and survival. This may contribute to off-target nuclear disruption and the induction of cellular senescence. In contrast, ICMT inhibitors act downstream of farnesylation and preserve the hydrophobic modification required for membrane interactions of other lamins, while selectively disrupting the production of progerin. Given that ICMT inhibition stimulates cellular proliferation whereas FTIs do not, these mechanistic differences suggest that ICMT inhibitors may represent a less disruptive and promising therapeutic strategy.

This research could be technically expanded by automating the solution exchange process using microfluidic systems to evaluate the faster-acting mechanisms occurring during and immediately after drug delivery^82–88^. Alternative labeling schemes involving nanobody arrays^89^ and fluorogenic dyes^90^ enable replenishable recruitment of multiple fluorophores to a single locus with improved signal and trajectory length, which could significantly extend tracking capabilities over longer periods of time. Switching to imaging modalities using thinner light sheets^87,88,91–93,93^ could also improve contrast, especially when imaging in the nuclear interior, where unbound fluorophores contribute to high background. Additionally, implementing point spread function engineering^87,88,93–96^ would extend tracking capabilities to 3D^92,97,98^, giving a more complete picture of telomere dynamics.

Taken together, our results improve our understanding of the effects of HGPS on chromatin dynamics, with potential implications for other genetic disorders linked to nuclear lamina mutations. Our quantitative study also provides a direct comparison of the efficacy of different therapeutic strategies on restoring healthy chromatin dynamics in HGPS cells. Beyond HGPS, this study offers critical insights into the fundamental processes of biological aging. Progerin is known to be produced in small amounts in aging cells of healthy individuals^28,99^, and nuclear defects are a hallmark of normal cellular senescence^99–101^. By investigating the link between lamina integrity and telomere dynamics, this research provides a deeper understanding of the markers of cellular aging.

## Methods

### Cell culture

Healthy and HGPS human fibroblasts (HGADFN167 and HGFDFN168, The Progeria Research Foundation) were cultured at 37°C and 5% CO_2_ in high glucose Dulbecco’s Modified Eagle’s Medium (DMEM) without L-glutamine (31053028, Gibco) and supplemented with 15% (v/v) fetal bovine serum (FBS) (A5669701, Gibco), 1% (v/v) penicillin-streptomycin (15140122, Gibco), and 1% GlutaMAX (35050061, Gibco).

293T cells (ATCC, CRL-3216) were cultured at 37°C and 5% CO_2_ in DMEM (ATCC 30-2002) supplemented with 10% (v/v) FBS (Gibco, A5669701) and 2 mM L-glutamine (ATCC 30-2214).

### Plasmid and lentivirus generation

A third-generation lentiviral transfer vector for expression of mSG-TRF2 was generated using standard molecular cloning techniques. First, amplicons of mStayGold and TRF2 containing compatible overhangs for Gibson assembly were generated by PCR (NEB, M0492S) from plasmids pcDNA3/F-tractin=mStayGold (Addgene #212019, a gift from Atsushi Miyawaki) and pCIBN-tagBFP-hTRF2 (Addgene #103803, a gift from Karsten Rippe), respectively. The two fragments were assembled into a PCR-linearized third-generation lentiviral transfer vector backbone after the Ubc promoter using NEBuilder HiFi Assembly (NEB, E2621S) following the manufacturer’s protocol, and 2 μL of the product was transformed into NEB Stable Competent E. coli (High Efficiency) cells (NEB, C3040H) following the manufacturer’s protocol. The plasmid was miniprepped (QIAGEN, 12123) and checked via full-plasmid sequencing (Plasmidsaurus). Third-generation lentivirus was produced via transfection of 293T cells (ATCC, CRL-3216) in a 6-well plate with 710 ng of plasmid pMDLG/pRRe (Addgene #12251, gift from Dr. Didier Trono), 455 ng of plasmid pMD2.G (Addgene #12259, gift from Dr. Didier Trono), 710 ng of plasmid pRSV-Rev (Addgene #12253, gift from Dr. Didier Trono), and 625 ng of the mSG-TRF2 transfer vector using Lipofectamine 3000 (ThermoFisher, L3000001) following the manufacturer’s protocols. Supernatant was harvested 24 and 48 h post transfection and concentrated using PEG-it viral precipitation kit (System Biosciences, LV810A-1) following the manufacturer’s protocols, aliquoted in PBS, and stored at −80°C.

### Drug treatment

HGPS fibroblasts (HGADFN167, The Progeria Research Foundation) were treated with either an FTI or an ICMT inhibitor to test the response of treatment on telomere dynamics. The FTI, Lonafarnib (SML1457-5MG, Sigma Aldrich), was diluted to 2 µM^36,68–70^ in HGPS media and used to treat cells for 2 days before transduction. The ICMT inhibitor C75^76^ (provided by the Bergö lab) was diluted to either 5 µM^65^ or 2 µM in HGPS media and used to treat the cells for 20 days or 10 days, respectively, before transduction. Control cells were treated with the vehicle (0.02 % (v/v) DMSO) in HGPS media for 2 days before transduction.

### Sample preparation and transduction

First, 8-well ibidi chambers (80807, ibidi) were coated with a 0.1% solution of fibronectin (FO895-1MG, Millipore Sigma) diluted 1:10 in Hanks’ Balanced Salt Solution (HBSS) (14-175-095, Fisher Scientific) and left to fully dry. Next, for transient mSG-TRF2 expression, cells were transduced with lentivirus 48 h before imaging. For this process, 40 μL of concentrated lentivirus was mixed with 35 μL of media containing 8 µg/mL polybrene (TR1003G, MilliporeSigma) before being added to the ibidi well. Then, 225 μL of cell suspension diluted to 5 x 10^4^ cells/mL in polybrene media was added to the well dropwise. The media was changed and replaced with regular cell media after 24 h. Finally, 1 h before imaging, 3 µM of SiR-DNA live-cell fluorescence nuclear stain (SC007, Cytoskeleton) diluted in cell media was added to the ibidi well to enable nuclear tracking to facilitate drift correction (Supplementary Fig. 6).

### Imaging procedure and settings

Cells were imaged on an inverted microscope (IX-83, Olympus) with an exposure time of 50 ms every 30 s for 45 min. The cells were kept in a live-cell incubator (Okolab, OKO-H301PI736ZR1/2S) and maintained at 37°C and 5% CO_2_ at 90% humidity during all experiments. A custom-built flat-field HILO illumination platform with a high-NA objective lens (UPLAPO100XOHR, 100X, NA 1.5, Olympus) was used as described elsewhere^102^. A 200 mW, 488 nm laser (1254728, Coherent) was used at 55 W/cm^2^ for illumination in the telomere channel for all datasets. A 1000 mW 647 nm laser (2RU-VFL-P-1000-647-FC/APC, MPB Communications) was used at 27 W/cm^2^ for illumination in the SiR-DNA-stained nuclear channel for all datasets. MicroManager was used for data acquisition to automate stage positioning and to synchronize the 488 nm channel and the 647 nm shutters. For every frame at each position, an 11-slice z-stack was acquired in 500 nm steps using MicroManager’s Z-stacks setting in the Multi-Dimensional Acquisition window. The maximum intensity projection of each frame was then acquired in post-processing to track telomeres across all z-planes concurrently. Between 10 – 12 cells were imaged per imaging session using MicroManager’s Multiple Positions (XY) setting in the Multi-Dimensional Acquisition window.

### Tracking and data analysis

For visualization of the HGPS nucleus in Fig. 1a, images of the nuclear DNA channel and the telomere-labeled channel were acquired separately and merged in post-processing where a rolling ball with a radius of 50 pixels was used for background subtraction on the telomere channel.

For telomere tracking in Figs. 1-3, first, two-channel image stacks of mSG-TRF2-transduced cells were cropped manually around the cell of interest to produce two separate image stacks: one including the cell with telomeres expressed, and one including the SiR-DNA-stained nuclear channel (Supplementary Fig. 6a,b). The telomeres were then localized with ThunderSTORM^103^ (Supplementary Fig. 6c) using the following parameters: filter = wavelet filter (B-Spline), order = 3, scale = 2.0, detection method = local maximum, peak intensity threshold = 2*std(Wave.F1), connectivity = 8-neighbourhood, estimation method = PSF: integrated gaussian, fitting radius = 3 pixels, sigma = 1.6 pixels, fitting method = weighted least squares, and spurious localizations were filtered out by filtering to include only localizations with uncertainties less than 25 nm. Duplicate localizations were removed by filtering out molecules that converged to the same position with a distance threshold of 120 nm.

The SiR-DNA-stained nuclear channel (Supplementary Fig. 6b) was used for center-of-mass-based drift correction. First, a 2-pixel radius Gaussian blur filter was applied in ImageJ via Process > Filters > Gaussian Blur to improve homogeneity of the overlayed nuclear mask (Supplementary Fig. 6d). Next, a binary mask was applied to the nucleus via Image > Adjust > Threshold (Supplementary Fig. 6e). Once the threshold was set, the center of mass of the nucleus in every frame across the image stack was calculated via Analyze > Set measurements > Center of mass and limit to threshold, then Analyze > Analyze particles > Show overlay masks (Supplementary Fig. 6f). Then, a custom MATLAB code was used to drift correct the telomere localizations and correct for cell movement over time by using the center of mass of the nucleus across the image stack.

The drift-corrected telomere localizations were then exported to trackpy.link^104^ for trajectory linking and analysis. Trajectories were linked using a maximum jump of 0.79 µm, a frame gap of 3 frames, a minimum track length of 60 frames, and a maximum lag of 15 frames. Then, time-averaged MSDs of individual trajectories were calculated using trackpy.ismd^104^ after particle linking and filtering to retain trajectories of fixed length (60 frames) to provide consistency for scan area analysis (Fig. 2). Scan areas of individual telomere trajectories were computed as the area of the 2D convex hull enclosing all positions of a given trajectory. Scan areas were then calculated per cell and pooled across replicates for each condition for statistical analysis (Fig. 2f). For visualization, tracks were colored according to their scan area, and mean scan areas across all cells for each replicate were reported (Supplementary Fig. 2).

For each trajectory, MSDs were calculated for the first 15 lags and were fit to Eq. 1 describing two-dimensional fractional Brownian motion, where D_eff_ is the effective diffusion coefficient, α is the anomalous exponent, and σ is the localization uncertainty. Model parameters were estimated by non-linear least-squares fitting in log-log space with parameter bounds as follows: D_eff_ ∈ [0.000001, 0.1] µm^2^/s^α^, α ∈ [0.01, 2], σ ∈ [0.001, 0.1] µm. Fits with α < 0.05 or α > 1.95 indicated poor fits and were excluded from further analysis.

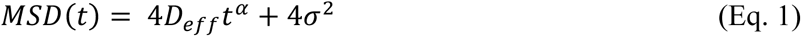

For visualization, MSD curves and violin plots were colored by condition (Fig. 3) and by replicate (Supplementary Fig. 4). Jump distances and angles were computed directly from linked trajectories via Eq. 2 and Eq. 3, respectively. In Eq. 2, 𝑟 is the 2D displacement vector, which represents the displacement in telomere position from one frame to the next. Therefore, ||𝑟|| is the magnitude of that displacement vector and represents the length of the telomere jump. In Eq. 3, consecutive jump angles were calculated between successive displacement vectors 𝑟_%_ and 𝑟_%&’_ and expressed in degrees spanning 0° to 180°. Steps with zero displacement were excluded from angle calculations. Jump distances were pooled per condition (Fig. 3d and Supplementary Fig. 5a) and were also calculated as per-cell median and pooled per condition for statistical analysis (Fig. 3e). Jump angles were pooled per condition (Fig. 3f and Supplementary Fig. 5b) and also calculated as per-cell medians and pooled per condition for statistical analysis (Fig. 3g).

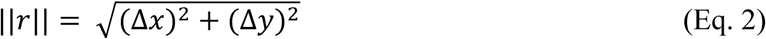

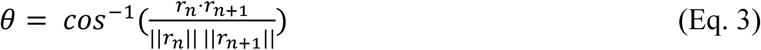

### Statistics and reproducibility

Tracking data was collected from three independent sample replicates for each condition. Each replicate received its own lentiviral transduction and was imaged on separate days. All data was checked for batch effects before pooling. Information on the number of trajectories and cells for each experiment can be found in Supplementary Tables 1-3. No statistical method was used to predetermine sample size throughout this work.

For statistical significance testing of the telomere tracking data in Fig. 2f, telomere scan areas were first calculated as per-cell mean values across replicates for all six conditions to avoid pseudo-replication with individual trajectories. All groups were then tested with a one-way ANOVA and all groups except the 5 μM C75-treated group were found to not have batch effects between replicates (control: *p* = 0.3 across replicates; HGPS: *p* = 0.5 across replicates; Lonafarnib-treated: *p* = 0.8 across replicates; 5 μM C75-treated: *p* = 0.005 across replicates; 2 μM C75-treated: *p* = 0.3 across replicates). Data was then pooled together per condition and a two-sided Mann–Whitney U tests was used for statistical significance testing, except in the case of the 5 μM C75-treated group which was tested with a linear mixed-effects model. All telomere trajectories across all conditions for each replicate are included in Supplementary Fig. 2 and full statistics including number of cells, mean values, and p values can be found in Supplementary Table 1.

For statistical significance testing of the telomere tracking data in Fig. 3b,c, diffusion coefficients and anomalous exponents were first calculated as per-cell median values. MSDs and violin plots for each technical replicate are shown in Supplementary Fig. 4. All groups were tested for batch effects with a one-way ANOVA and all groups except the Lonafarnib-treated group and 5 μM C75-treated groups did not show batch effects (control: D_eff_ *p* = 0.3 across replicates and α *p* = 0.7 across replicates; HGPS: D_eff_ *p* = 0.6 across replicates and α *p* = 0.4 across replicates; Lonafarnib-treated: D_eff_ *p* = 0.02 across replicates and α *p* = 0.06 across replicates; 5 μM C75-treated: D_eff_ *p* = 0.3 across replicates and α *p* = 0.01 across replicates; 2 μM C75-treated: D_eff_ *p* = 0.7 across replicates and α *p* = 0.6 across replicates). In the case of groups with batch effects (Lonafarnib-treated, D_eff_ and 5 μM C75-treated, α) a linear mixed-effects model with condition as a mixed effect and replicate as a random effect was used to gauge statistical significance. All other groups were pooled together per-condition and a two-sided Mann–Whitney U test was used for statistical significance. Full statistics including number of cells per condition, median D_eff_ values, median α values, and all p values can be found in Supplementary Table 2.

For statistical significance testing of the telomere tracking data in Fig. 3e,g, jump distances and jump angles were first calculated as per-cell median values and were tested for batch effects between replicates with a Kruskal-Wallis test before being pooled across the three replicate groups per condition. Kruskal-Wallis test per condition results showed batch effects for the control, HGPS, and 5 μM C75-treated groups (control: jump distance *p* = 0.08 across replicates, jump angle *p* = 0.03 across replicates; HGPS: jump distance *p* = 0.03 across replicates, jump angle *p* = 0.6 across replicates; Lonafarnib-treated: jump distance *p* = 0.8 across replicates, jump angle *p* = 0.4 across replicates; 5 μM C75-treated: jump distance *p* = 0.002 across replicates, jump angle *p* = 0.03 across replicates; 2 μM C75-treated: jump distance *p* = 0.5, jump angle *p* = 0.1). Statistical comparison between conditions was performed using a linear mixed-effects model with condition (HGPS vs other groups and controls vs other groups) as a mixed effect and replicate as a random effect. Statistical comparisons between all conditions including number of cells, number of jumps, mean jump distances, and p values can be found in Supplementary Table 3.

## Data availability

Localizations and trajectories generated and analyzed during the current study and source data that can be used for generating the graphs in this work are provided on GitHub [https://github.com/Gustavsson-Lab/chromtrack/tree/main].

## Code availability

Telomeres were localized using the open-source FIJI plugin ThunderSTORM^103^ [https://github.com/zitmen/thunderstorm/releases/tag/v1.3/]. Localizations were linked into a trajectory using trackpy.link^104^ [https://soft-matter.github.io/trackpy/dev/generated/trackpy.link.html]. MSDs were calculated using trackpy.imsd^104^ [https://soft-matter.github.io/trackpy/dev/generated/trackpy.motion.imsd.html#trackpy.motion.imsd]. Fits were performed and plots were generated using custom code in Python with common packages, available on GitHub [https://github.com/Gustavsson-Lab/Progeria-Telomere-Tracking---Source-Data-and-Analysis]. A custom drift correction code was written in MATLAB using the center of mass of the nucleus and is available on Github [https://github.com/Gustavsson-Lab/Progeria-Telomere-Tracking---Source-Data-and-Analysis].

## Acknowledgements

This work was supported by the National Institute of General Medical Sciences of the National Institutes of Health grant R35GM155365 and startup funds from the Cancer Prevention and Research Institute of Texas grant RR200025 to AKG. The authors thank The Progeria Research Foundation for donation of the human fibroblast cell lines used in this work (HGADFN167 and HGFDFN168).

## Author contributions

G.G. transduced the cells, performed the experiments, analyzed the data, and wrote the manuscript. A.R. generated the plasmids and lentivirus used for transduction, the tracking analysis pipeline, and contributed to the manuscript. X.Y. provided the ICMT inhibitor C75 used in this work. M.B. contributed to data interpretation and feedback on the manuscript. A.-K.G. conceived the idea, contributed to the manuscript, and supervised the research.

## Competing interests

The authors declare that they have no competing interests.

## Notes

### Competing Interest Statement

The authors have declared no competing interest.

