## Supplementary Information for "Increased Telomere Mobility in Progeria is Restored by Isoprenylcysteine Carboxyl Methyltransferase Inhibition"

### Supplementary Figures

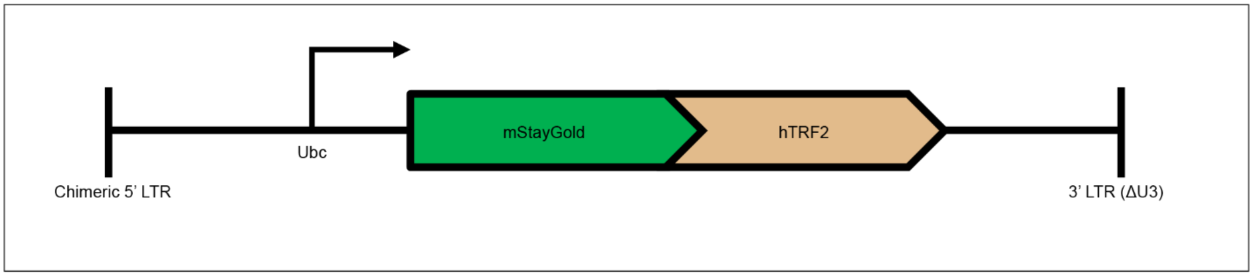


**Supplementary Fig. 1. Simplified plasmid schematic for the telomere label.** The fluorescent protein mStayGold is fused onto the N-terminus of human TRF2 (hTRF2), separated by a short peptide linker. The construct is driven by the Ubc promoter. This cassette is placed in a 3$^{rd}$ generation lentiviral transfer vector enabling production of lentiviral particles used to transduce cells for telomere tracking experiments.

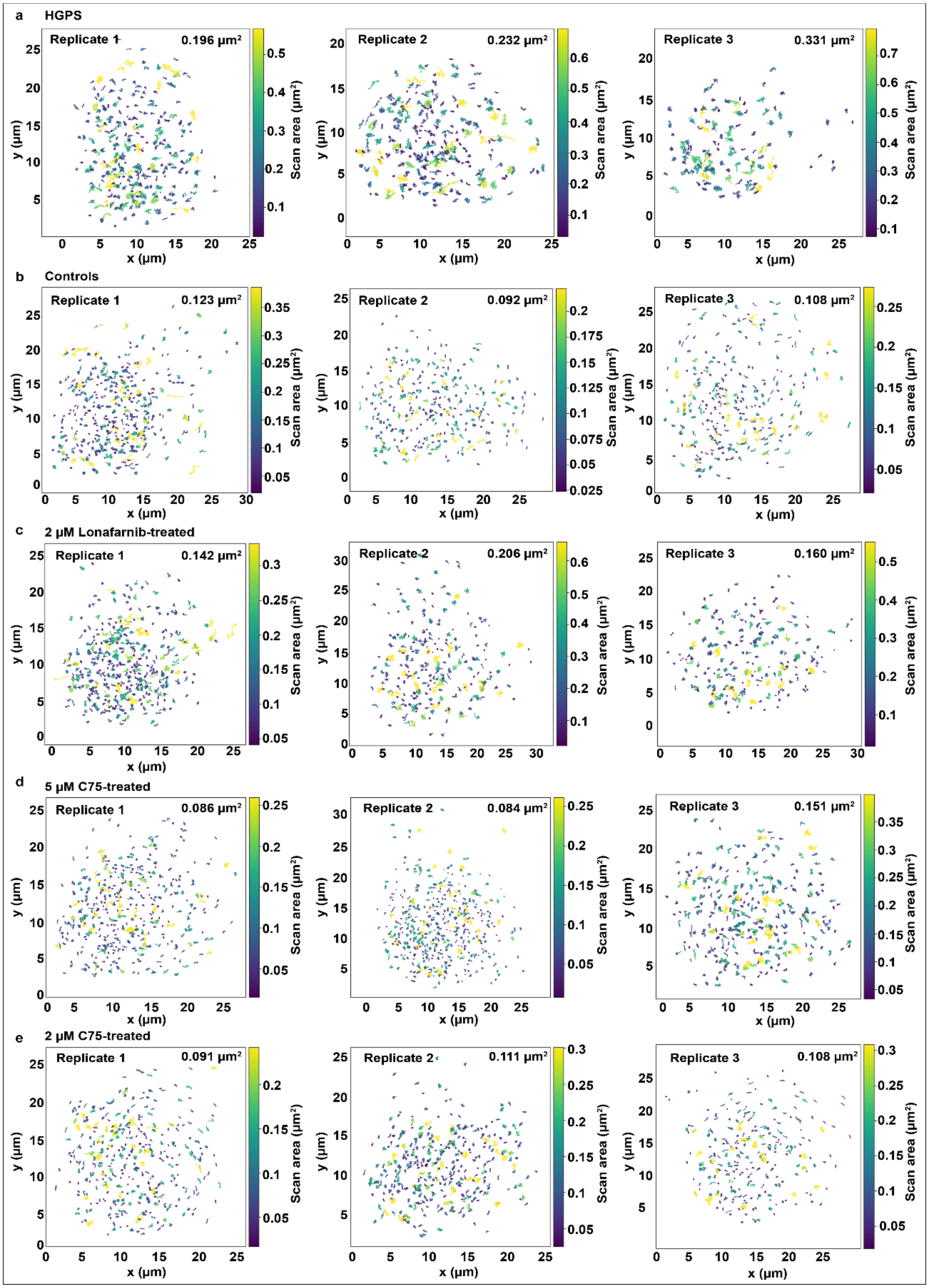


**Supplementary Fig. 2. Telomere scan areas for condition replicates.** Telomere trajectories in **a** HGPS cells, **b** control cells, **c** 2 μM Lonafarnib-treated HGPS cells, **d** 5 μM C75-treated HGPS cells, and **e** 2 μM

C75-treated-HGPS cells for replicates 1, 2, and 3 colored by scan area. Mean scan areas per replicate are shown in the top right corner of each box.

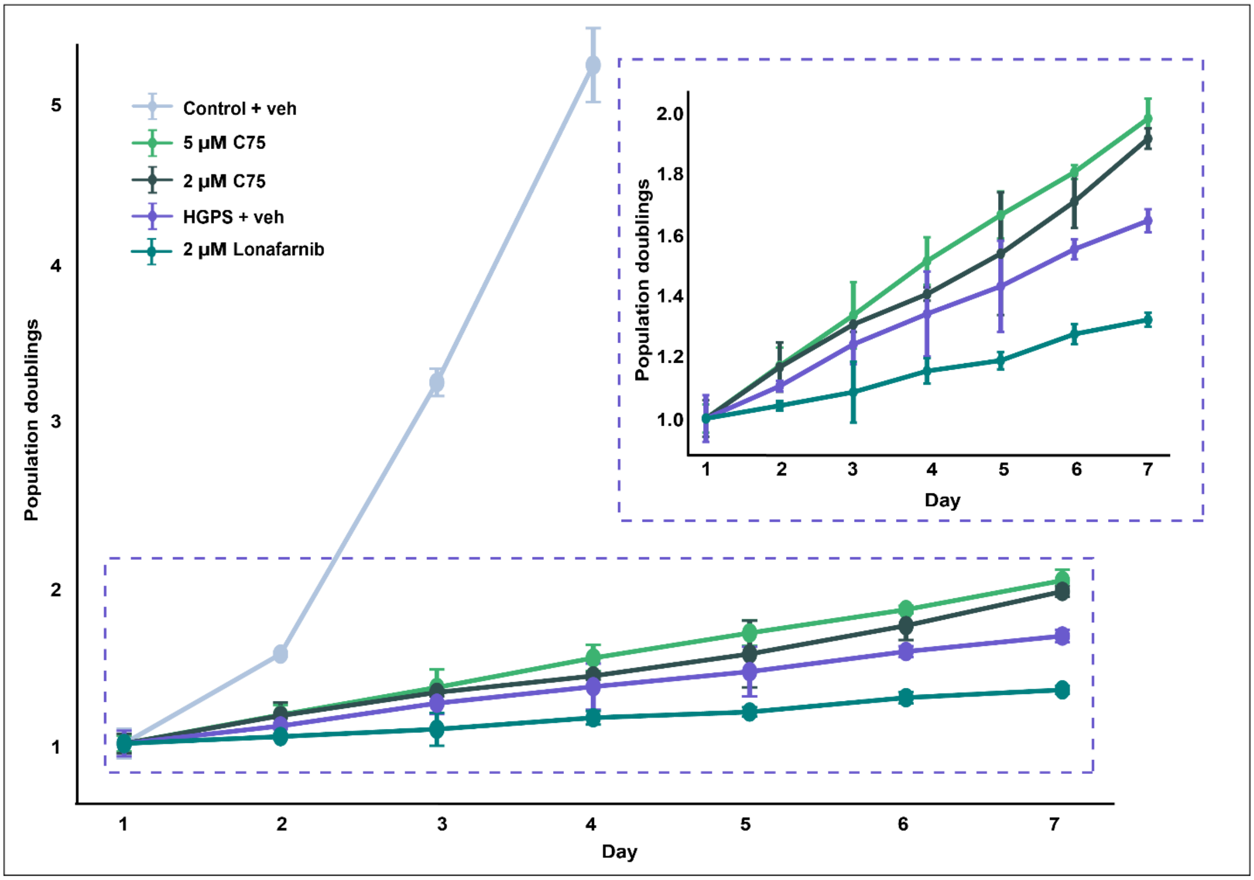


**Supplementary Fig. 3. Growth curves for control, HGPS, and drug-treated cell lines.** Growth curves showing the mean ± SEM of the cell population doubling per day over 7 days. Three different fields of view (FOV) per day were imaged per condition and cells were counted using ImageJ's MultiPoint tool. Cell numbers were normalized to the day 1 mean for each condition. Healthy control cell counting was stopped after 4 days due to cells reaching confluency. Inset shows a zoomed-in portion of the area in the dotted region, without the control cell line, to gauge difference between HGPS treatments.

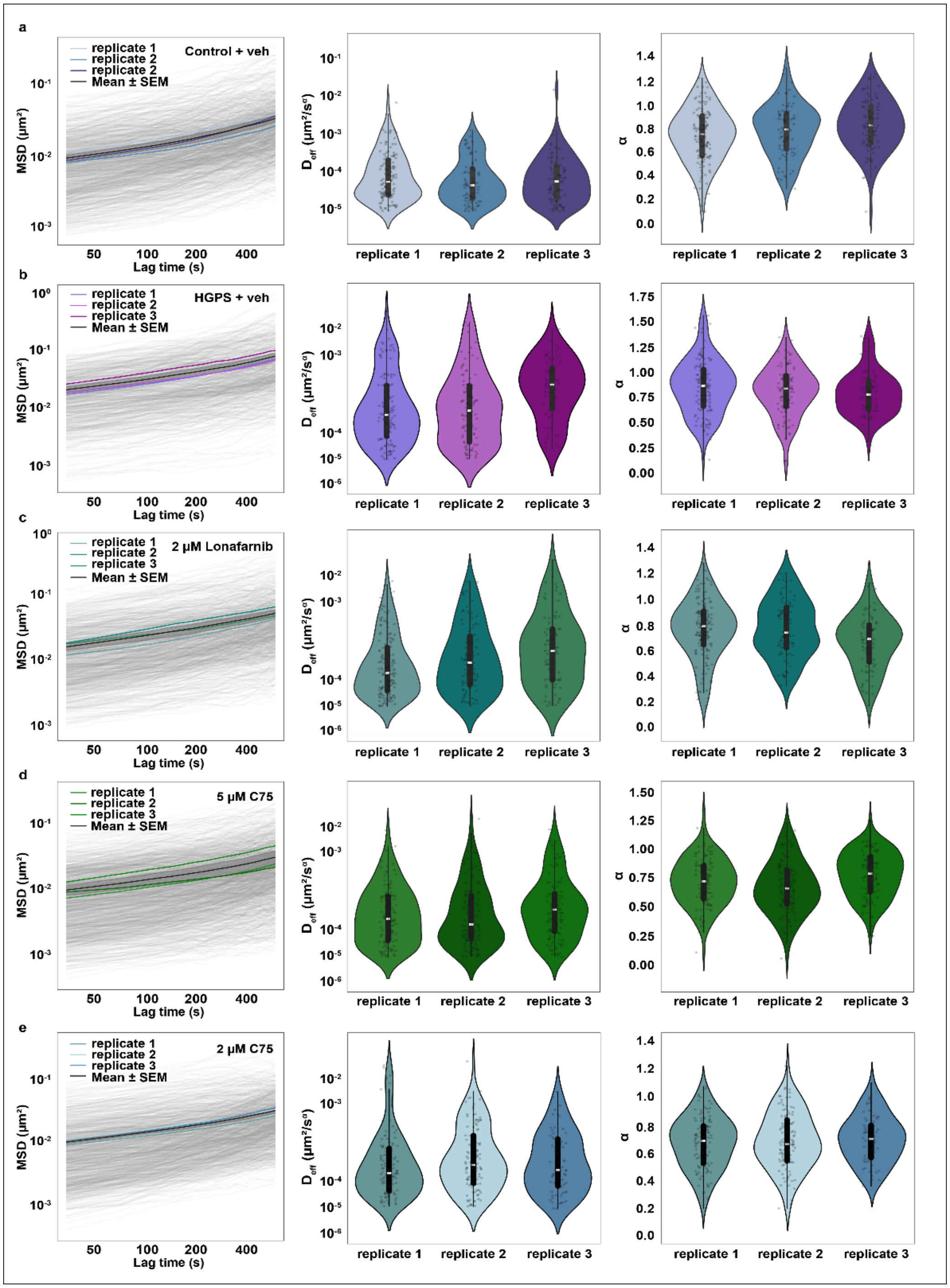


**Supplementary Fig. 4. MSD analysis and violin plots for condition replicates.** MSD curves (left), violin plots for the effective diffusion coefficient ($D_{eff}$) (middle), and violin plots for the anomalous exponent (α) (right) for three technical replicates of telomere trajectories in **a** healthy control cells, **b** HGPS cells, **c** HGPS cells treated with 2 μM Lonafarnib for 2 days, **d** HGPS cells treated with 5 μM C75 for 20 days, and **e** HGPS cells treated with 2 μM C75 for 10 days. All MSD curves are shown colored by individual replicate means, and with the mean ± SEM of the three replicates shown in black. All violin plots include a black rectangle representing the interquartile range from the 25$^{th}$ to 75$^{th}$ percentile, and a white line at the center representing the median.

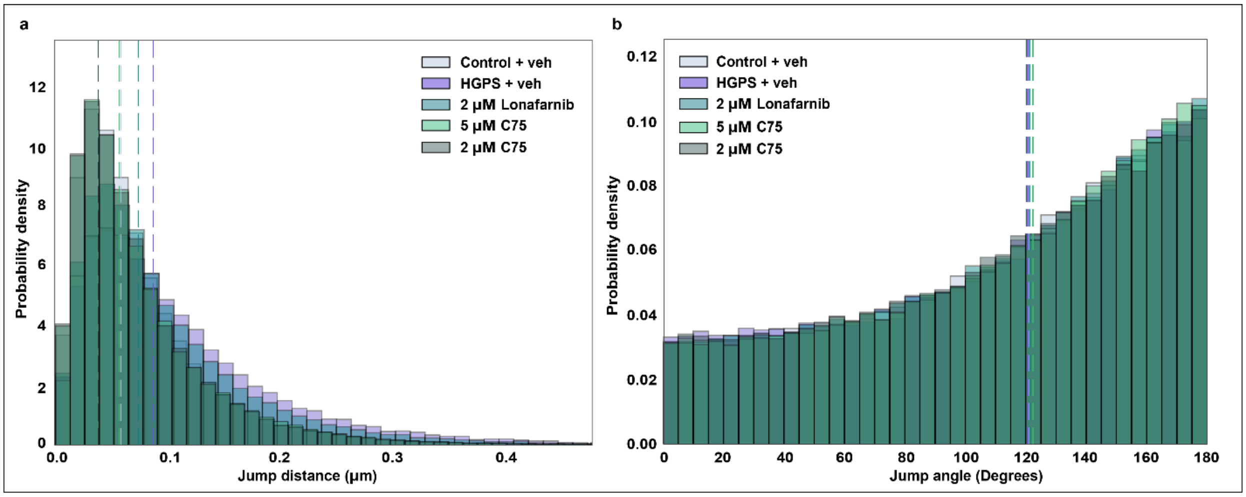


**Supplementary Fig. 5. Telomere jump distance and jump angle histograms across conditions.** **a** Histograms comparing telomere jump distances across all conditions. All individual jumps from all tracked telomeres were pooled across three technical replicates acquired on separate days for each condition (see Supplementary Table 2 for all statistics). Dashed vertical lines indicate the median jump distance for each group which were 0.058 µm for controls, 0.086 µm for HGPS, 0.074 µm for Lonafarnib-treated, 0.056 µm for 5 µM C75-treated, and 0.056 µm for 2 µM C75-treated. **b** Histograms of consecutive jump angles for telomeres across all conditions, calculated as the angle between successive displacement vectors along individual trajectories and demonstrating anti-persistent motion (180-degree bias). All angles were pooled across the three technical replicates per condition (see Supplementary Table 2 for all statistics). Dashed vertical lines indicate the median jump angle for each group which were 121.2° for controls, 120.7° for HGPS, 121.3° for 2 µM Lonafarnib-treated, 122.4° for 5 µM C75-treated, and 120.2° for 2 µM C75-treated.

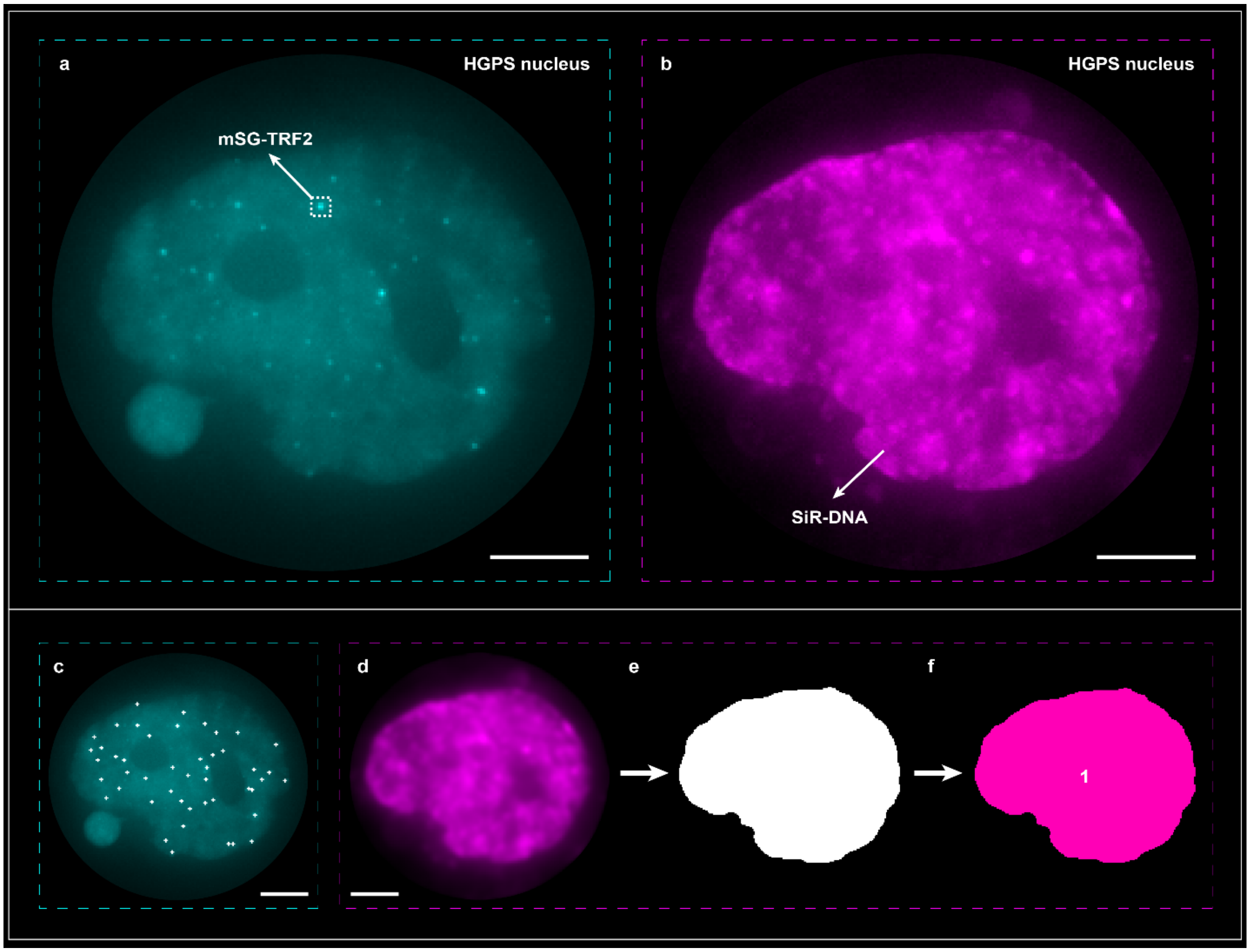


**Supplementary Fig. 6. Telomere drift correction scheme with nuclear center of mass.** Two-color imaging of **a** telomeres shown in cyan and **b** nuclear DNA labeled with SiR-DNA stain shown in magenta, imaged in separate color channels. Scale bars 5 µm. **c** Post-processing of telomere data from **a**, where telomeres are localized in ImageJ's Thunderstorm plugin, shown as crosses colored in white. **d,e,f** Post-processing of nuclear DNA channel data, where first (**d**), a 2-pixel radius Gaussian blur filter is applied to improve the homogeneity of the binary mask applied in **e**. **f** Next, the center of mass of the nuclear mask is calculated and labeled for each frame, which is then used for drift correction of the telomeres in **c**. Scale bars 5 µm.

**Supplementary Table 1. Telomere scan area statistics for Fig. 2f.** Statistics for telomere diffusion data shown in Fig. 2f. Data was tested for significance with either a two-sided Mann–Whitney U test (M.W.) or a linear mixed effects model (L.M.M.).

| Condition | *n* (cells) | Mean scan area ($\mu m^2$) | Scan area *p* values (vs HGPS) | Scan area *p* values (vs controls) |
|---|---|---|---|---|
| **Control (WT)** | 45 | 0.128 | $2.71 \times 10^{-8}$ (M.W.) | - |
| **HGPS** | 43 | 0.318 | - | $2.71 \times 10^{-8}$ (M.W.) |
| **2 µM Lonafarnib** | 43 | 0.157 | $1.84 \times 10^{-5}$ (M.W.) | 0.001 (M.W.) |
| **5 µM C75** | 38 | 0.098 | $3.6 \times 10^{-7}$ (L.M.M.) | 0.03 (L.M.M.) |
| **2 µM C75** | 32 | 0.104 | $3.25 \times 10^{-9}$ (M.W.) | 0.6 (M.W.) |

**Supplementary Table 2. Telomere dynamics statistics across conditions for Fig. 3b,c.** Statistics for telomere diffusion data shown in Fig. 3b,c. Data was tested for significance with either a two-sided Mann–Whitney U test (M.W.) or a linear mixed effects model (L.M.M.).

| **Condition** | ***n* (cells)** | **Median $D_{eff}$ ($\mu m^2/s^{\alpha}$)** | **$D_{eff}$ *p* values (vs HGPS)** | **$D_{eff}$ *p* values (vs controls)** | **Median $\alpha$** | **$\alpha$ *p* values (vs HGPS)** | **$\alpha$ *p* values (vs controls)** |
|---|---|---|---|---|---|---|---|
| **Control (WT)** | 45 | $1.66 \times 10^{-4}$ | 0.007 (M.W) | - | 0.56 | 1.0 (M.W.) | - |
| **HGPS** | 43 | $3.89 \times 10^{-4}$ | - | 0.007 (M.W.) | 0.62 | - | 1.0 (M.W.) |
| **2 µM Lonafarnib** | 43 | $3.27 \times 10^{-4}$ | 1.0 (L.M.M) | 0.03 (L.M.M.) | 0.49 | 0.3 (M.W.) | 0.3 (M.W.) |
| **5 µM C75** | 38 | $2.05 \times 10^{-4}$ | 0.03 (M.W.) | 0.5 (M.W.) | 0.47 | 0.02 (L.M.M.) | 0.04 (L.M.M.) |
| **2 µM C75** | 32 | $1.62 \times 10^{-4}$ | 0.005 (M.W.) | 0.9 (M.W.) | 0.53 | 0.7 (M.W.) | 0.5 (M.W.) |

**Supplementary Table 3. Telomere jump distance and jump angle statistics for Fig. 3e,g.** Statistics for telomere jump distance and jump angle data shown in Fig. 3e,g. Data was tested for significance with a linear mixed effects model (L.M.M.).

| Condition | *n* (cells), *n* (jumps), *n* (angles) | Mean jump distance (µm) | Jump distance *p* values (vs HGPS) | Jump distance *p* values (vs controls) | Mean jump angle (°) | Jump angle *p* values (vs HGPS) | Jump angle *p* values (vs controls) |
|---|---|---|---|---|---|---|---|
| **Control (WT)** | 45, 58,000, 57,955 | 0.064 | $1.14 \times 10^{-11}$ (L.M.M.) | - | 120.0 | 0.1 (L.M.M.) | - |
| **HGPS** | 43, 39,440, 39,397 | 0.101 | - | $1.14 \times 10^{-11}$ (L.M.M.) | 117.0 | - | 0.1 (L.M.M.) |
| **2 µM Lonafarnib** | 43, 54,142, 52,099 | 0.076 | $7.47 \times 10^{-6}$ (L.M.M.) | $3.82 \times 10^{-3}$ (L.M.M.) | 121.6 | 0.02 (L.M.M.) | 0.1 (L.M.M.) |
| **5 µM C75** | 38, 70,760, 70,722 | 0.059 | $7.29 \times 10^{-17}$ (L.M.M.) | 0.2 (L.M.M.) | 122.8 | 0.006 (L.M.M.) | 0.03 (L.M.M.) |
| **2 µM C75** | 32, 57,826, 57,794 | 0.060 | $1.06 \times 10^{-12}$ (L.M.M.) | 0.4 (L.M.M.) | 120.3 | 0.1 (L.M.M.) | 0.7 (L.M.M.) |
